# Chromatin compartments define distinct modes of action and biological roles of the pioneer Pax7

**DOI:** 10.64898/2026.08.21.746263

**Authors:** Arthur Gouhier, Juliette Harris, Vincent Lapointe-Roberge, Alexandre Marcil, Aurelio Balsalobre, Jacques Drouin

## Abstract

Pioneer transcription factors direct cell differentiation by remodeling closed chromatin to deploy new enhancers. It remains unclear how the chromatin context shapes their action. During pituitary development, the pioneer PAX7 opens thousands of enhancers and specifies the melanotrope cell fate. We report that PAX7 acts in both the A (active) and B (inactive) genomic compartments, engaging two distinct closed states: H1-enriched chromatin in A, and H3K9me2- and lamin-rich chromatin in B. Whereas A-compartment enhancers open in a single, cell-division-independent step, B-compartment opening requires cell division and triggers domain-wide lamin dissociation coupled with B-to-A compartment shift. The two modes involve distinct biological outputs: A-compartment action modulates broadly expressed genes, whereas B-compartment action drives de novo expression of cell-type-specific developmental genes central to melanotrope identity.

## Introduction

Pioneer transcription factors (PF) specify cell fates by binding their target DNA sequences within closed chromatin and initiating its remodeling to deploy new enhancer repertoires that drive lineage-specific gene expression (*1, 2*). Recent structural and biochemical studies have clarified how pioneers engage nucleosomal DNA and recruit the chromatin-remodeling cofactors that complete enhancer opening (*3-5*). However, binding nucleosomal DNA is not sufficient for productive pioneer action: many PF binding sites are occupied without being opened (*6-8*), suggesting that features of the surrounding chromatin environment determine whether remodeling proceeds (*9*).

Closed chromatin environments are heterogeneous and vary with regards to linker histone H1 levels, lamina association and histone modifications, notably H3K27me3-enriched domains repressed by the Polycomb complex and H3K9me3-enriched domains that recruit HP1 proteins (*10, 11*). These environments differ in compaction mechanism, 3D organization within the nucleus, and partition into the active A and inactive B compartments and finer sub-compartments They also differ in their permissiveness to PF binding: H3K9me3-marked constitutive heterochromatin is largely refractory to PF action (*1, 6*), whereas lamina-associated chromatin can be permissive (*13*), and its reorganization in aging and disease redirects binding of the PF FoxA2 (*14, 15*). In addition to opening individual enhancers, PFs were associated with larger-scale chromatin reorganization, including topologically associating domain (TAD) opening and restructuring and enhancer-promoter loop formation (*16-18*). However, the relationship between local pioneer engagement at individual enhancers and large-scale compartment reorganization, and whether compartment context shapes pioneer mechanism and biological output, has not been systematically addressed.

The PF PAX7 specifies the melanotrope fate in the pituitary intermediate lobe (*8, 19*). Using the corticotrope AtT-20 cell line that is reprogramed towards a melanotrope-like identity by *Pax7* ectopic expression (Fig. 1A), we previously showed that PAX7 rapidly primes its target enhancers but requires passage through cell division for full chromatin opening, with the cell-cycle-dependent step involving H3K9me2 demethylation and dissociation from the nuclear lamina (*8, 13*). By integrating data from Micro-C, chromatin markers, gene expression and in vivo *Pax7* loss-of-function in mouse pituitary, we now report that PAX7 acts as a pioneer in both A and B compartments through two mechanistically distinct modes. By contrast to a direct single-step activation for enhancers in the A compartment, PAX7-initiated chromatin opening in the B compartment requires cell division and results in domain-wide lamin dissociation coupled with B-to-A compartment shift. The two modes are associated with functionally distinct biological outputs: A-compartment action modulates broadly expressed genes encoding general biological functions such as signaling and secretory activity, whereas B-compartment action drives de novo expression of cell-type-specific developmental genes central to melanotrope identity.

**Fig. 1.**
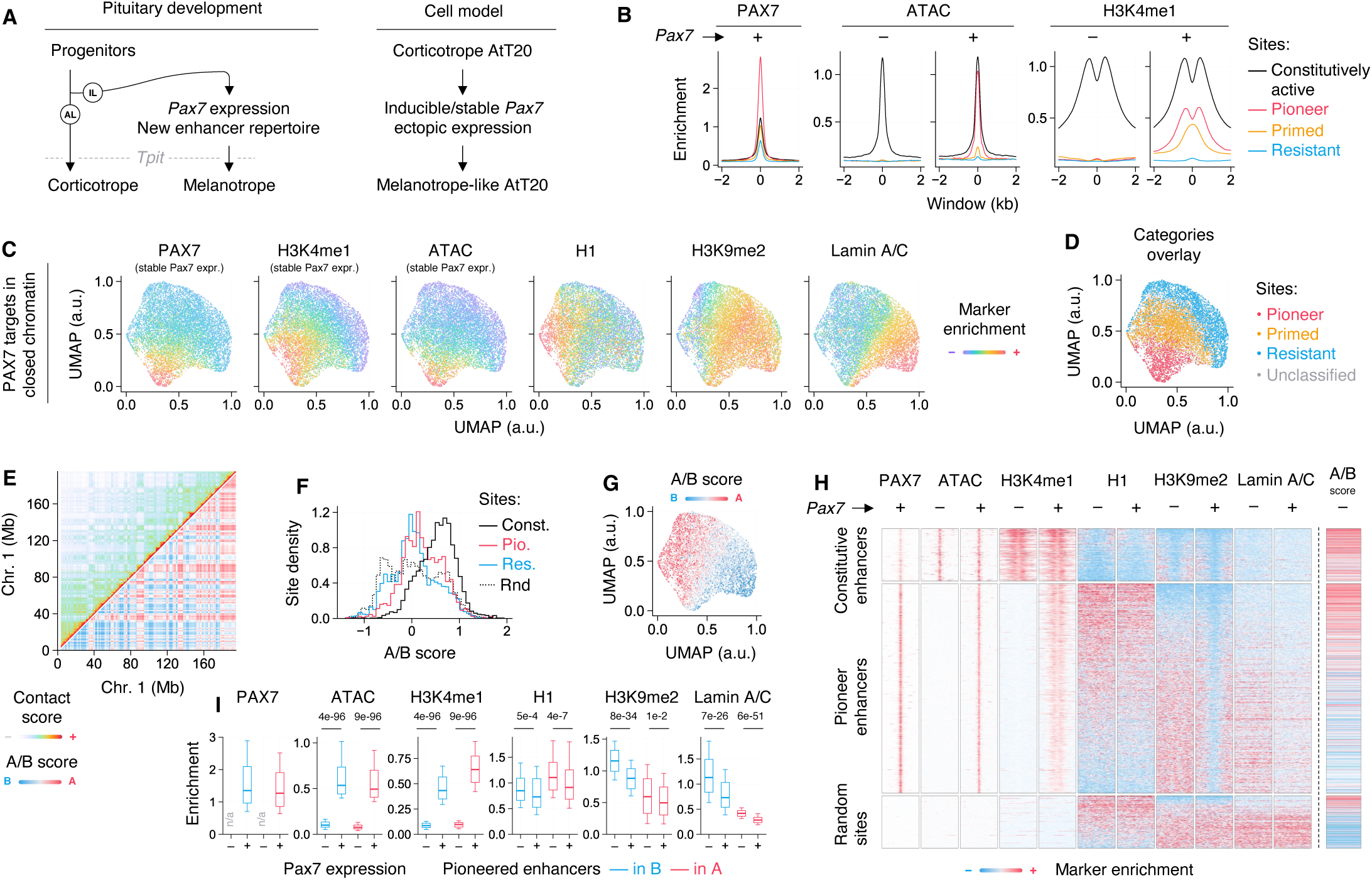
PAX7 acts as pioneer factor in both A/B genomic compartments. (**A**) Schematic representation of PAX7 role in specifying intermediate lobe (IL) melanotrope cell fate by contrast the corticotrope cell fate in the anterior lobe (AL). Both lineages express the same hormone precursor gene, proopiomelanocortin (POMC), but not the final hormone. Their terminal differentiation is controlled by the Tbox transcription factor TPIT. (**B**) Mean profile of the indicated markers with or without stable *Pax7* expression at multiple subsets of PAX7-targeted sites in closed chromatin: constitutively active enhancers (n = 3,984), pioneer (n = 1,152), primed (n = 2,523) and resistant sites (n = 2,761). (**C**) UMAP generated with the indicated markers at PAX7 targets in closed chromatin (n = 9,321). (**D**) Categorization of PAX7 targets in closed chromatin from (B) overlayed on UMAP from (C). (**E**) Micro-C contact map at resolution of 100 kb of AtT20 chromosome 1 in absence of *Pax7* expression (top left). Corresponding A/B compartment identity (bottom right). (**F**) Histogram of A/B score distribution at the specified categories of PAX7-targeted sites. (**G**) A/B score at PAX7 targets in closed chromatin, UMAP from (C). (**H**) Heatmaps of the indicated markers at multiple subsets of PAX7-targeted sites with or without stable *Pax7* expression. Constitutive and random sites subsampled. Sites ordered by the ratio of initial H1 over H3K9me2 signal. 4 kb window centered on PAX7 recruitment. (**I**) Quantification of the specified markers with or without stable *Pax7* expression at the ¼ pioneer enhancers with strongest B association or A association. P-values computed from two-sided Mann-Whitney U tests; boxplots represent the 10-25-50-75-90th percentiles.

## Results

### PAX7 targets in closed chromatin are heterogeneous

To understand how the initial chromatin environment influences PAX7 pioneering activity, we first assessed DNA accessibility using the assay for transposase-accessible chromatin (ATAC) and the deposition of the activating histone H3 lysine 4 monomethylation mark (H3K4me1) by chromatin immunoprecipitation and sequencing (ChIP-seq) at PAX7 recruitment sites (fig. S1A, table S1). Of 28,504 PAX7 binding sites (excluding promoters and insulators, where PAX7 was shown to have no effect on chromatin structure (*8*)), 9,321 sites are located in closed chromatin, characterized by an absence of DNA accessibility (ATAC) and H3K4me1 deposition before *Pax7* expression. We further classified these closed targets with respect to their response to PAX7 recruitment: Pioneer enhancers (n = 1,152), where PAX7 leads to chromatin opening and H3K4me1 gain; Primed enhancers (n = 2,523), with deposition of H3K4me1 but weak chromatin opening; and Resistant sites (n = 2,761), where no change is observed (Fig. 1B, fig. S1B). Compared to constitutively active enhancers, closed targets are enriched for linker histone H1 as well as for the repressive H3K9me2 mark. Consistent with the recognition of H3K9me2 by lamina-associated proteins (*20*), closed targets also show a stronger association with the nuclear lamina, as assessed by Lamin A/C ChIP-seq (fig. S1A,C). At Pioneer enhancers, PAX7 leads to a local loss of histone H1, best revealed by the higher resolution of ChIP-qPCR (*13*), and of H3K9me2, as well as a broader lamin dissociation (fig. S1C,D). This is consistent with our published model in which lamin dissociation is required for PAX7-initiated enhancer activation However, UMAP representation of chromatin markers at closed PAX7 targets revealed that these sites segregate into two distinct populations: one initially enriched for linker histone H1 and the other enriched for H3K9me2 with stronger lamin association (Fig. 1C). Notably, there is no apparent correlation between these two chromatin states and the responsiveness to PAX7, as both Pioneer enhancers and Resistant sites are similarly distributed between these states (Fig. 1D). This observation raised the question of whether these chromatin states reflect a broader organizational feature of the genome.

### PAX7 acts as pioneer factor in both genomic compartments

We hypothesized that the two chromatin states observed at closed PAX7 targets might correspond to the higher-order organization of the genome into A (active) and B (inactive) compartments. To test this, we performed Micro-C (genome-wide chromosome conformation capture) (*21*) in AtT-20 cells before (Fig. 1E) and after *Pax7* expression (fig. S2A). Principal component analysis of chromosome-wide contact maps identified topologically associating domains (TADs) and resolved A and B genomic compartments (fig. S2B). The A compartment shows a general enrichment for the activating H3K4me1 mark but also for linker histone H1, whereas the B compartment is enriched for the repressive H3K9me2 mark and displays stronger lamin association (fig. S2C). This is consistent with the typical classification of A and B compartments as transcriptionally active and inactive, respectively (*12*). The histone H1 antibody used here is marketed as pan-H1 but the A-compartment-enriched distribution we observe is consistent with the H1.4 variant being preferentially detected (*22-24*). We next assessed PAX7 binding sites in relation to genomic compartments. Whereas constitutively active enhancers are mostly located in the A compartment, both Pioneer and Resistant targets are distributed across both compartments (Fig. 1F). Importantly, A and B compartments recapitulate the two distinct chromatin states previously observed at closed PAX7 targets, as shown by UMAP (Fig. 1G) and heatmap (Fig. 1H, fig. S2D) representations. This correspondence validates the interpretation that the two populations identified in our initial UMAP analysis (Fig. 1C) reflect compartment identity. Consistently, histone H1 loss occurs predominantly at Pioneer enhancers in the A compartment, while H3K9me2 loss and lamin dissociation occur at targets in the B compartment (Fig. 1I). Notably, genome compartmentalization does not affect PAX7 final recruitment nor the gain in DNA accessibility (ATAC) and enhancer activation (H3K4me1, Fig. 1I). Thus, despite engaging two fundamentally distinct chromatin environments, PAX7 achieves comparable levels of binding and enhancer activation in both compartments, suggesting that it employs distinct mechanisms to reach the same functional outcome.

### Sub-compartments are differentially permissive to PAX7 action

Although both Pioneer and Resistant targets share a similar distribution across A and B compartments, we hypothesized that finer sub-compartments characterized by distinct chromatin properties (*12, 25*) might distinguish permissive from non-permissive environments for PAX7 pioneering activity. To identify such sub-compartments, we performed principal component analysis on trans contacts between even and odd chromosomes, thereby isolating the compartment signal from cis-contact bias (Fig. 2A,B). The first component recapitulates the overall A/B compartmentalization, with the activating H3K4me1 and H3K4me3 marks colocalizing with the A compartment (Fig. 2C). However, repressive histone modifications and lamin association show notable variation in their distribution across the B compartment (Fig. 2C). K-means clustering of this landscape resolved five distinct clusters (Fig. 2D, table S2). Clusters I and II correspond to very active (I) and active (II) regions within the A compartment, showing the strongest deposition of H3K4me1 and H3K4me3 (Fig. 2E). Clusters III, IV and V are associated with the B compartment and display significant levels of both H3K9me2 and lamin association (Fig. 2E). Beyond this shared signature, cluster IV is distinguished by high deposition of H3K27me3 (Fig. 2E), reflecting Polycomb-mediated repression, as supported by the recruitment of PRC1, PRC2 and deposition of H2AK119Ub (fig. S3A,B). Notably, PRC2 recruitment is higher in clusters III and V relatively to cluster IV (fig. S3A, B), suggesting a structural role independent of its representative mark (*26*). Cluster V is characterized by high H3K9me3 and the strongest lamin association, consistent with constitutive heterochromatin (Fig. 2E).

**Fig. 2.**
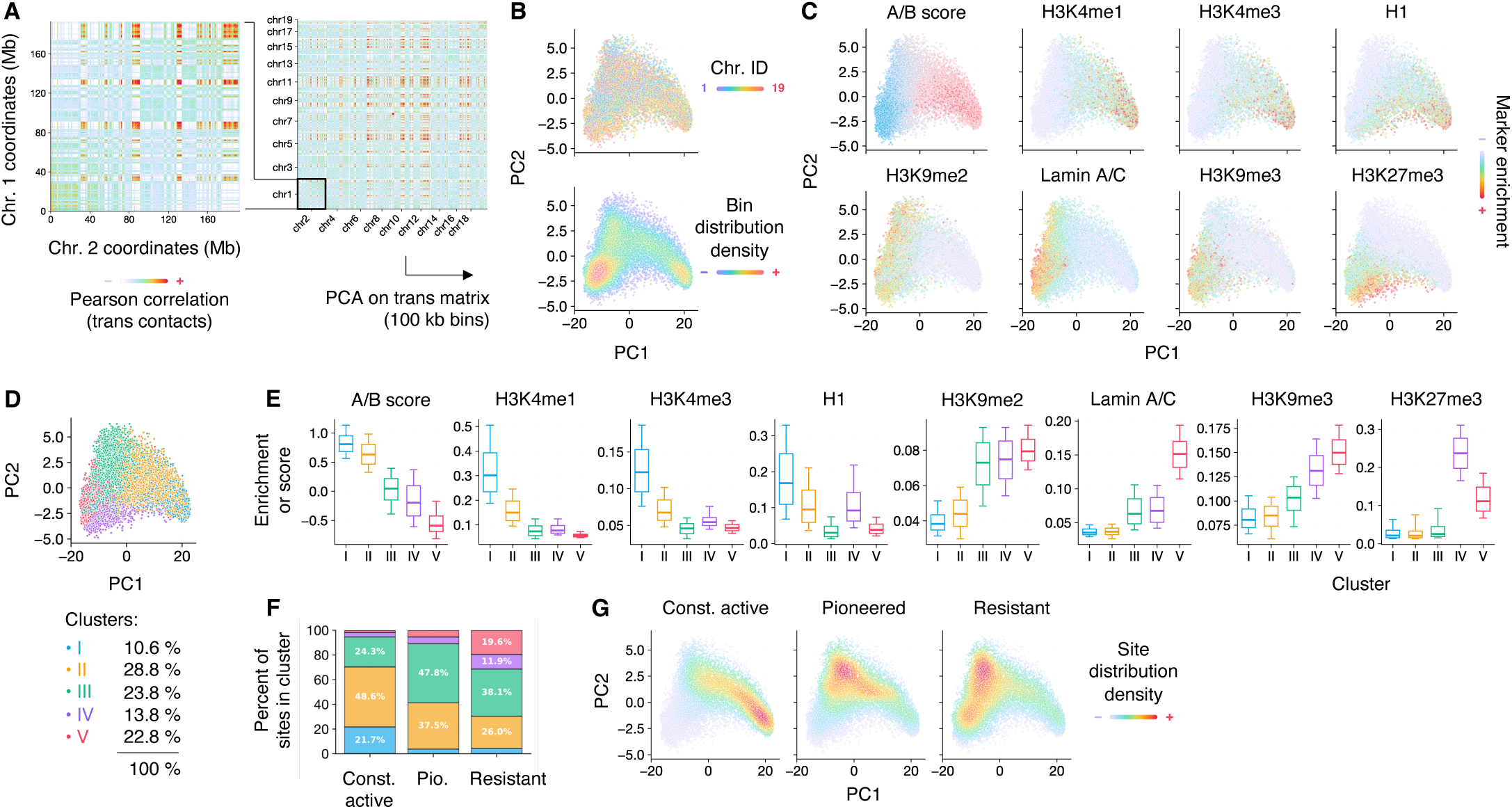
Sub-compartments are differentially permissive to PAX7 action. (**A**) Genome-wide Pearson correlation matrix of KR-normalized Micro-C trans contact scores at a resolution of 100 kb across all odd vs. even chromosomes. (**B**) PCA on matrix from (A) at a resolution of 100 kb, with bin chromosome ID or distribution density color-coded. (**C**) Average value of the indicated marker per bin from (B). (**D**) K-means cluster association using chromatin landscape from (C) of each bin from (B). (**E**) Quantification of the specified markers without *Pax7* expression at the clusters from (D). All pairwise comparisons are statistically significant. (**F**) Distribution of PAX7-targeted sites of the specified categories across clusters from (D). (**G**) Distribution density of PAX7-targeted sites of the specified categories overlaid on PCA from (B). P-values computed from two-sided Mann-Whitney U tests; boxplots represent the 10-25-50-75-90th percentiles.

Notably, Resistant sites are the only closed-chromatin PAX7 targets present in clusters IV and V (Fig. 2F,G, fig. S3C), suggesting that Polycomb-repressed and constitutively heterochromatic environments are refractory to PAX7 pioneer action. By contrast, Pioneer enhancers are found almost exclusively in clusters II and III (Fig. 2F,G). Thus, while PAX7 can pioneer enhancers across the full range of chromatin states spanning from H1-rich (cluster II, A compartment) to H3K9me2/lamin-rich (cluster III, B compartment), it appears unable to act in environments dominated by Polycomb-mediated repression or H3K9me3-enriched constitutive heterochromatin. These may define the boundaries of the chromatin landscape that is permissive to PAX7 pioneering activity.

### Cell division is required for PAX7-initiated chromatin opening in the B compartment

We previously showed that PAX7 pioneer action requires passage through cell division for lamin dissociation and chromatin opening (*13*). Having now established that Pioneer enhancers reside in two distinct compartments with markedly different levels of lamin association, we sought to assess whether the requirement for cell division is linked to the initial compartment. While PAX7 achieves comparable enhancer activation in both compartments in dividing cells (Fig. 1I), we asked whether this outcome depends on cell division in both cases. To do so, we arrested AtT-20 cells in G1 for 12 h using the G1-arrest agent mimosine and assessed the state of Pioneer enhancers after 48 h of activation of an inducible ER-PAX7 chimera (Fig. 3A). Both the inducible ER-PAX7 and mimosine systems were previously validated (*8, 13*).

**Fig. 3.**
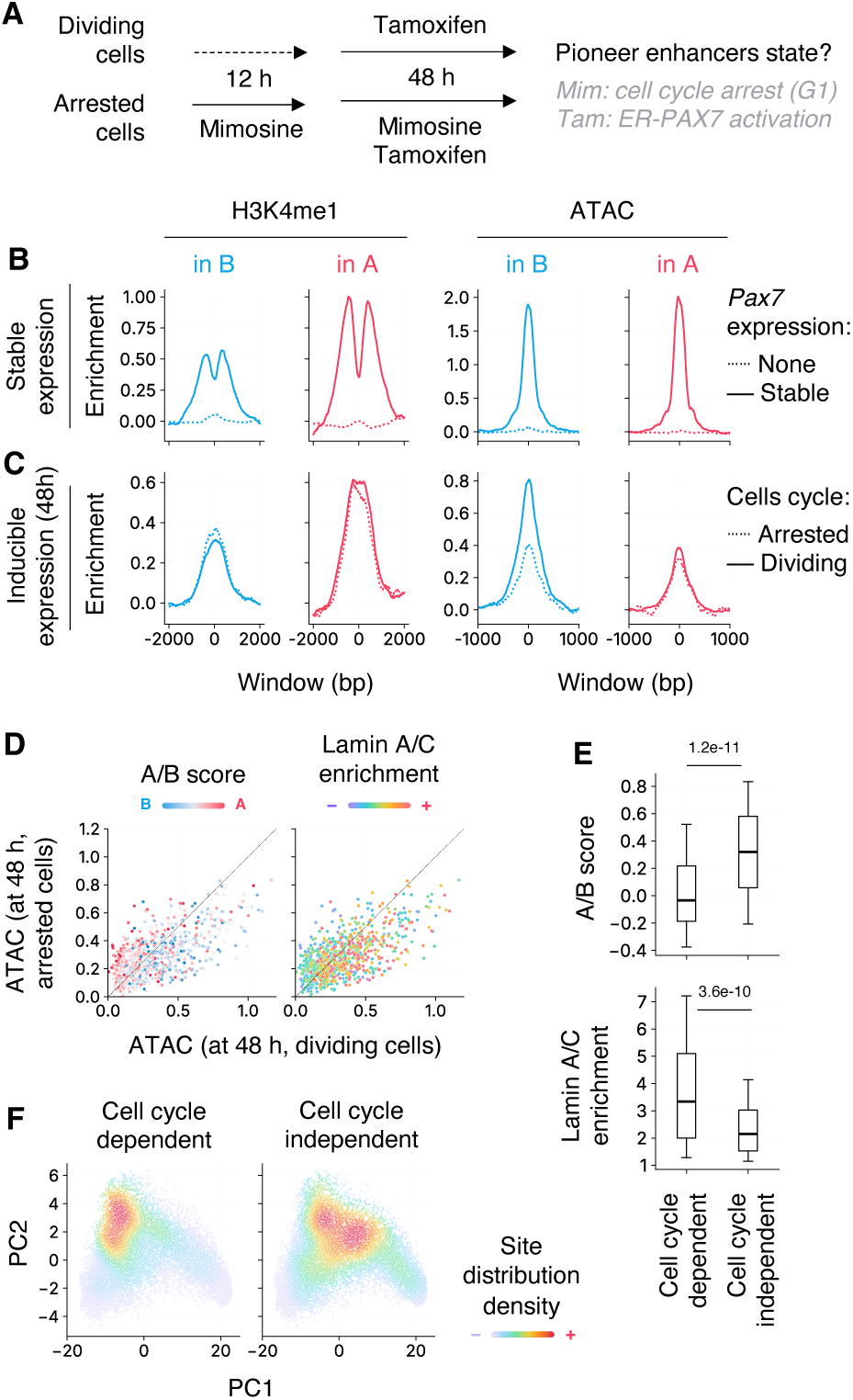
Cell division is required for PAX7-initiated chromatin opening in the B compartment. (**A**) Schematic representation of the experimental design to assess PAX7 action in cell-cycle-arrested vs. dividing cells. (**B**) Mean profile of H3K4me1 deposition and DNA accessibility (ATAC) at Pioneer enhancers with or without stable *Pax7* expression. (**C**) Mean profile of H3K4me1 deposition and DNA accessibility (ATAC) at Pioneer enhancers in cell-cycle-arrested vs. dividing cells after 48 h of ER-PAX7 activation. (**D**) DNA accessibility (ATAC) enhancers in cell-cycle-arrested vs. dividing cells after 48 h of ER-PAX7 activation correlated with initial A/B score and lamin A/C association. (**E**) Quantification of the initial A/B score and lamin A/C association at the ¼ Pioneer enhancers with strongest dependence or independence on cell division for DNA accessibility gain between 0 and 48 h of ER-PAX7 activation. (**F**) Distribution density of PAX7-targeted sites from (E) overlaid on Micro-C PCA from Fig. 2B. P-values computed from two-sided Mann-Whitney U tests; boxplots represent the 10-25-50-75-90th percentiles.

At 48 h, we observed no difference in H3K4me1 deposition between Pioneer enhancers in the A and B compartments (Fig. 3B,C), consistent with the cell division-independent monomodal deposition of H3K4me1 (priming) that follows PAX7 recruitment (*13*). By contrast, we observed a reduction in DNA accessibility gains specifically for targets in the B compartment, but not in the A compartment (Fig. 3B,C). It should be noted that Pioneer enhancers were defined here using stable *Pax7* expression, and we previously showed that PAX7 pioneering can require multiple days, particularly for targets with high initial H1 levels (*13*), which accounts for the overall modest DNA accessibility gains observed at Pioneer targets in the A compartment at this time point (Fig. 3C). Despite this limitation, we observed a clear correlation between initial B compartment and lamin association, and the loss of DNA accessibility between arrested and dividing cells (Fig. 3D). In addition, when we split Pioneer enhancers based on their dependence on cell division (defined as whether cell cycle arrest leads to a reduction in DNA accessibility gain at 48 h) the cell division-dependent group was significantly more associated with both the B compartment and lamin A/C (Fig. 3E). Consistently, cell division-dependent Pioneer enhancers map to sub-compartment cluster III (B compartment), whereas cell division-independent enhancers are preferentially located in cluster II (A compartment) (Fig. 3F). The presence of cell division-independent enhancers in cluster III likely reflects sites that have not yet undergone chromatin opening at this time point, rather than a true independence from cell division. Altogether, these results indicate that cell division is required for PAX7-initiated chromatin opening in the B compartment, while no such dependence is observed in the A compartment.

### PAX7 causes domain-wide B-to-A compartment shift

Given the established link between lamina association and B compartment identity (*27*), and our observation that PAX7 causes broad lamin dissociation at targets in the B compartment (Fig. 1H,I), we hypothesized that PAX7-induced lamin loss might trigger a shift in compartment identity. We first examined this at the *Pcsk2* locus, which encodes the proprotein convertase central to melanotrope identity and is differentially accessible in vivo between melanotrope and corticotrope cells (*16*). At this locus, PAX7 causes lamin dissociation that extends to the TAD boundaries, and this is associated with a TAD-wide opening and B-to-A compartment shift (Fig. 4A,B,C).

**Fig. 4.**
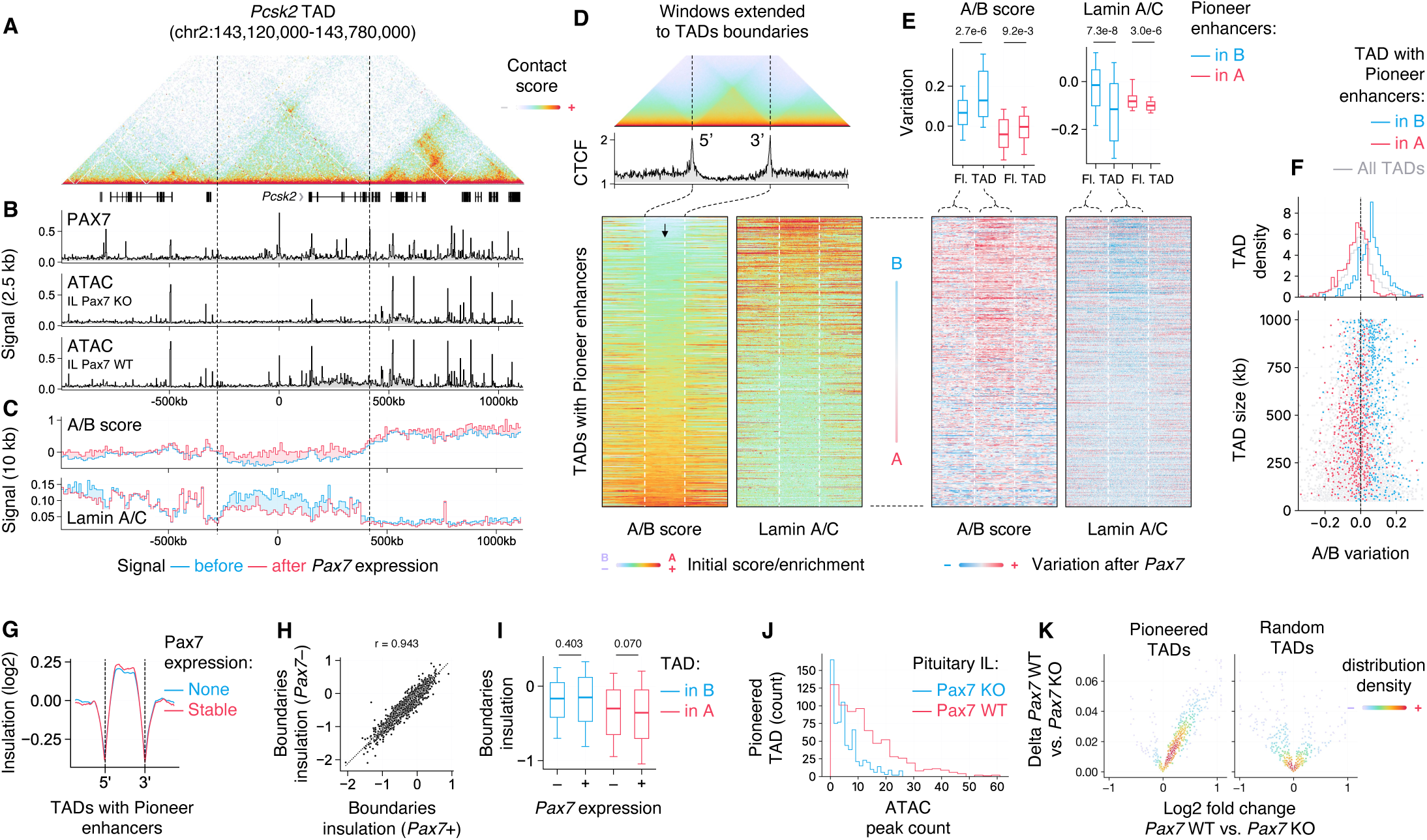
PAX7 causes domain-wide B-to-A compartment shift. (**A**) Micro-C contact map and GENECODE M25 gene annotation at the TAD harboring the *Pcsk2* gene, centered on PAX7-pioneered enhancer. (**B**) PAX7 recruitment in AtT-20 cells and ATAC-seq signal in *Pax7* WT vs. *Pax7* KO pituitary intermediate lobe. (**C**) A/B score and lamin A/C association with or without stable *Pax7* expression in AtT-20 cells. (**D**) Average Micro-C contact map and average CTCF ChIP-seq profile before *Pax7* expression at unique and unnested TADs harboring Pioneer enhancers (top). Heatmaps of initial A/B score and lamin A/C association at these TADs, ordered by average A/B score (bottom). (**E**) Heatmaps of A/B score and lamin A/C association variation before and after *Pax7* stable expression at TADs from (D). Distribution of the average variation within the TADs or the 200 kb flanking regions (Fl.) at the ¼ Pioneer enhancers with strongest B or A association is indicated on top. (**F**) Scatter plot of average TAD-wide A/B score variation at all TADs genome-wide vs. TAD size (bottom) and associated distribution histogram (top). TADs harboring one of the ¼ Pioneer enhancers with strongest B or A association are indicated in color. (**G**) Mean insulation score along the TADs and their flanking regions from (D) before and after stable *Pax7* expression, computed as the log2 mean off-diagonal contact frequency within a sliding diamond window. (**H**) Insulation score at individual TAD boundaries before vs. after stable *Pax7* expression (r = Pearson correlation coefficient). (**I**) Insulation score at TAD boundaries before and after stable Pax7 expression, separated by their initial compartment. (**J**) Distribution of ATAC-seq peaks in TADs from (D) in *Pax7* WT vs. *Pax7* KO pituitary intermediate lobe. (**K**) Volcano plot of average ATAC-seq signal (below ATAC-seq peaks) in TADs from (D) and random TADs in *Pax7* WT vs. *Pax7* KO pituitary intermediate lobe. P-values computed from two-sided Mann-Whitney U tests; boxplots represent the 10-25-50-75-90th percentiles.

To determine whether this domain-wide shift is a general feature of PAX7 pioneering in the B compartment, we expanded our analysis to all Pioneer enhancers. We performed TAD calling (table S3) and assigned each pioneered enhancer to a single TAD; in the case of nested TADs, we selected the TAD exhibiting the largest difference in compartment shift relative to its flanking regions (Fig. 4D). Lamin association and compartment identity are consistent within TADs, and the correlation between lamin association and B compartment identity observed locally at Pioneer enhancers (Fig. 1H,I) is recapitulated at the TAD level (Fig. 4E). Pioneer enhancers in the B compartment exhibit TAD-wide lamin dissociation and B-to-A compartment shift following *Pax7* expression, while no such changes are observed at Pioneer enhancers in the A compartment (Fig. 4E). This distinction is also confirmed when assessing the A/B variation at all TADs containing a Pioneer enhancer, irrespective of their size and nested configuration (Fig. 4F). PAX7 recruitment to closed chromatin is not sufficient per se to initiate a B-to-A compartment shift, as TADs containing only Primed enhancers, where PAX7 does not lead to H3K9me2 loss and enhancer activation (*13*)), do not undergo this shift (fig. S4A,B). Notably, the lamin dissociation and B-to-A compartment shift occurring after *Pax7* expression are restricted to within the boundaries of the affected TADs, as both are significantly stronger within the TAD relative to flanking regions (Fig. 4E) and the TADs boundaries are not affected by PAX7 action (Fig. 4G-I). This suggests that TADs act as structural units that delimit the domain-wide chromatin reorganization initiated by PAX7 in the B compartment.

To assess PAX7 domain-wide action in vivo, we compared DNA accessibility at the TAD level in wild-type and *Pax7* knockout intermediate lobe pituitary cells (*16*). TADs identified as Pioneer-containing in AtT-20 cells show a marked increase in both the number of ATAC-seq peaks (Fig. 4J) and the overall DNA accessibility signal in wild-type relative to *Pax7* knockout tissue (Fig. 4K), confirming that TAD-wide remodeling is a genuine property of PAX7 action in the lineage in which it physiologically acts.

### PAX7 implements two distinct classes of biological processes

Having established that PAX7 employs distinct modes of action and domain-wide regulation depending on compartment identity, we next asked whether the transcriptional consequences of PAX7 action also differ between the A and B compartments. To do so, we analyzed RNA-seq data in AtT-20 cells before and after *Pax7* expression and identified 1,129 genes regulated by PAX7 (Fig. 5A). Of these, 60% reside in the A compartment and 40% in the B compartment, with a majority of genes being upregulated in both compartments (Fig. 5B,C). We next performed gene ontology (GO) analysis separately on genes regulated in each compartment (table S4). Strikingly, while PAX7-regulated genes in the A compartment are involved in general cellular functions, regulated genes in the B compartment are distinctly enriched for terms related to development and differentiation (Fig. 5D). This distinction holds irrespective of whether the genes are up or downregulated (fig. S5). Furthermore, while regulated genes in the A compartment show quantitative changes in expression, indicating modulation of already active genes, upregulated genes in the B compartment show de novo expression (Fig. 5E).

**Fig. 5.**
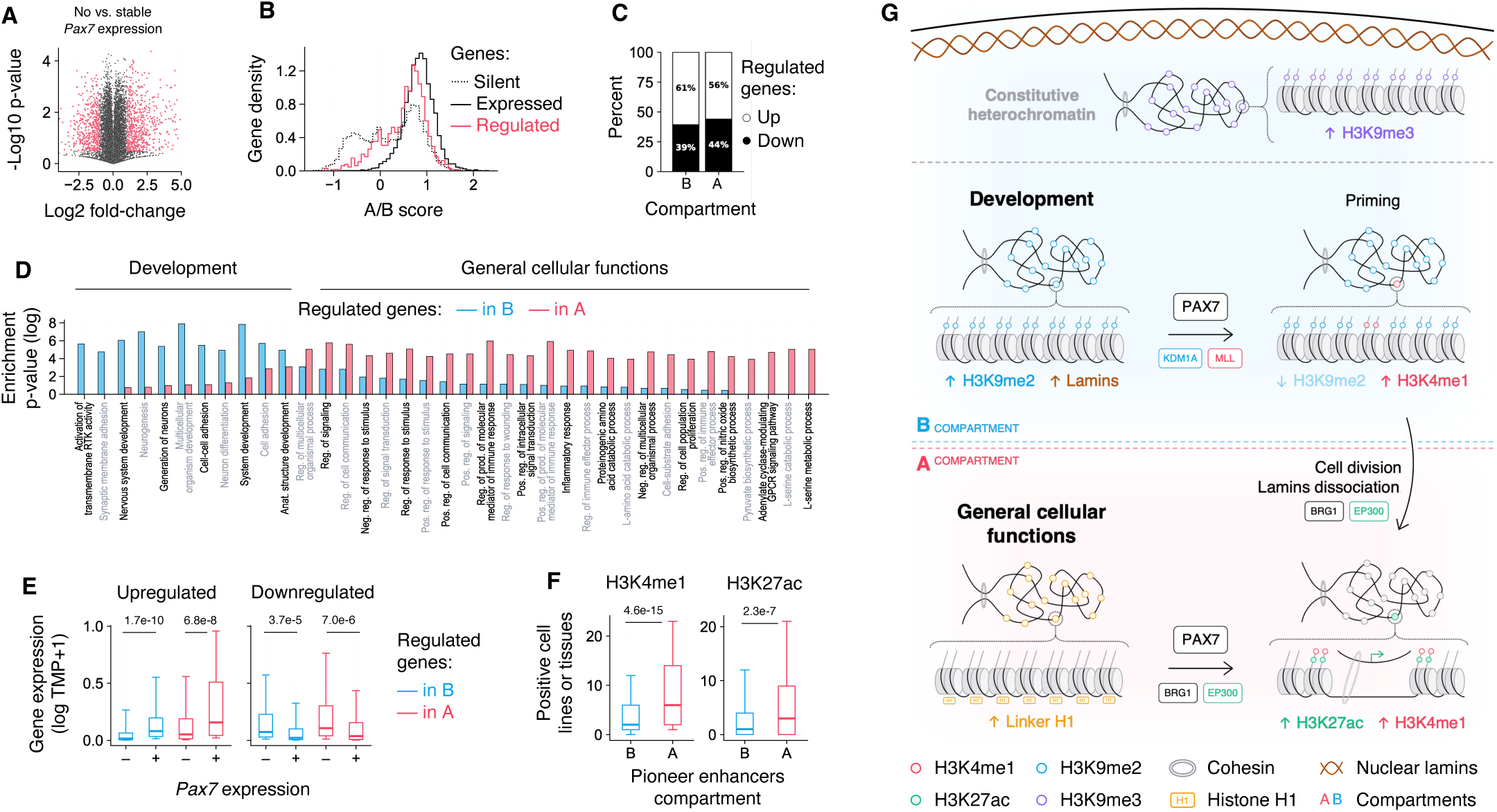
PAX7 implements two distinct classes of biological processes. (**A**) Differential gene expression of genes with or without stable *Pax7* expression. Genes above selection thresholds (absolute fold-change > 2 and p-value < 1e-5) are indicated in red (up = 620, down = 509). (**B**) A/B score distribution at expressed (TPM > 1), silent (TPM < 0.1) and PAX7-regulated promoters. (**C**) Distribution of gene expression variation between ¼ regulated genes with strongest B or A association. (**D**) Gene ontology (GO, biological process) analysis of PAX7-regulated genes in A (red) vs. B (blue) compartment. Terms ordered by the difference in A vs. B enrichment. Grayed terms for visualization purpose only. (**E**) Distribution of gene expression (transcript per million, TPM) with or without stable *Pax7* expression at ¼ regulated genes with strongest B or A association, split by their up or downregulation by PAX7. (F) Number of cell lines or tissues exhibiting H3K4me1 or H3K27ac deposition for the ¼ Pioneer enhancers with strongest B or A association. (**G**) Model of Pax7 pioneer action in the A and B compartments. P-values computed from two-sided Mann-Whitney U tests; boxplots represent the 10-25-50-75-90th percentiles.

The de novo activation of B compartment genes and their enrichment for developmental functions suggested that the associated enhancers are cell specific. To assess the cell-type specificity of PAX7-pioneered enhancers, we leveraged publicly available ChIP-seq peak-calling data from ChIP-Atlas (*28*), encompassing 2,008 H3K4me1 and 4,536 H3K27ac samples across 237 and 368 cell lines or tissues, respectively (table S5). For each pioneered enhancer, we determined whether it is positively marked in at least one sample per cell type. Pioneered enhancers in the B compartment are active in significantly fewer cell types than those in the A compartment (Fig. 5F), consistent with their association with cell-type-specific developmental genes.

Taken together, these results reveal that PAX7 action in the A compartment primarily modulates broadly expressed genes involved in general cellular functions, whereas its action in the B compartment activates previously silent, cell-type-specific genes that are central to the establishment of the new melanotrope cell fate.

## Discussion

This work shows that the pioneer factor PAX7 engages closed chromatin through two compartment-defined modes that converge on comparable enhancer activation but differ in kinetics, in their requirement for cell division, in the scale of associated chromatin reorganization, and in the biological function of the genes they regulate. This dual-mode framework integrates previously disparate observations on pioneer factors within a single organizing principle and identifies genomic compartmentalization not only as a downstream consequence of gene activity but as an active determinant of pioneer mechanism.

In the B compartment, PAX7 initially primes its target enhancers through the KDM1A-dependent loss of H3K9me2 and the MLL-associated gain of H3K4me1. It is only after passage through cell division that lamin association is lost, and the SWI-SNF remodelling complex and general coactivator EP300 are recruited, leading to nucleosome displacement and enhancer activation (*13*). PAX7 targets in the B compartment are present in domains enriched in H3K9me2, with high lamin association. Its action results in TAD-wide lamin dissociation and B-to-A compartment shift, as well as a marked increase in DNA accessibility across the TAD. Lamin dissociation and compartment shift are dependent on the local H3K9me2 loss caused by PAX7 at targeted enhancers: H3K9me2 mediates lamin reassociation following cell division (*20*), and inhibition of the KDM1A H3K9me2 demethylase prevents both PAX7-initiated H3K9me2 loss and lamin dissociation (*13*). Further, TADs encompassing only Primed enhancers where PAX7 recruitment leads to H3K4me1 deposition but not to H3K9me2 loss, do not exhibit such domain-wide changes. How the localized action of PAX7 at Pioneer enhancers results in domain-wide changes is unclear. Attractions between heterochromatic loci, whose structure is here altered by PAX7 action, have been found to be crucial for established compartmentalization (*27*). PAX7 also leads to cohesin recruitment following cell division (*13*) and to the formation of new genomic loops between loci (Harris et al. In submission). The combined changes in chromatin state and contact map inside TADs may thereby drive the domain-wide compartment shift.

This contrasts with PAX7 action in the A compartment, where no dependence on cell division nor domain-wide change in genomic compartmentalization is observed, consistent with low lamin association. PAX7 targets in the A compartment exhibit however markedly higher linker histone H1 levels. H1 promotes chromatin compaction genome-wide (*29, 30*) and pioneer factors have been shown to displace H1 when engaging closed chromatin (*13, 31, 32*). It imposes nevertheless a barrier to PAX7 action as its levels inversely correlate with the speed of PAX7 recruitment at Pioneer enhancers (*13*) and its loss facilitates re-activation of targets following PAX7 re-expression (Harris et al. In submission). Although multiple H1 variants exist and are associated with distinct nuclear localization (H1.2, H1.3 and H1.5 at the nuclear periphery whereas H1.4 is present throughout the nucleus) (*22, 23*), it is the variant H1.4 that is likely measured here according to its compartment distribution and that acts as a strong chromatin condenser (*33*) and limits PAX7 recruitment. Taken together, this supports that histone-H1-mediated chromatin compaction and H3K9me2/lamin-enriched chromatin environments impose distinct barriers to pioneer factor recruitment and action. Cell division is therefore not intrinsic to pioneering per se but is required specifically to license the B-to-A compartment shift that accompanies opening in the lamina-associated environment. This distinction may explain why replication dependence has appeared variable across pioneer systems (*7, 13, 34*): the requirement may depend on the compartment of the targets examined rather than reflecting a uniform property of the factor.

Sub-compartment dissection further defines the boundaries of pioneer action. Within the B compartment, PAX7 pioneers enhancers in cluster III (a lamin-associated, H3K9me2-enriched environment) but is refractory in cluster IV (Polycomb-repressed, H3K27me3-enriched) and cluster V (constitutive heterochromatin and H3K9me3-enriched). H3K9me3-enriched chromatin was shown to be refractory to other pioneers (*6, 7*). Here, H3K9me3 and H3K27me3 do not prevent PAX7 recruitment but rather its action. What distinguishes the permissive from the refractory clusters may be the removability of their defining mark: the H3K9me2 of cluster III is removed by PAX7-recruited KDM1A during priming (*13*), whereas H3K27me3 of cluster IV and H3K9me3 of cluster V are regulated by other demethylases and mechanisms (*35, 36*). The transition from H3K9me2 (cluster III) to H3K9me3 (cluster V) thus appears to mark a fundamental threshold separating facultative from constitutive heterochromatin in the context of cell-fate specification. In any case, chromatin permissive to PAX7 action in the B compartment is devoid of H3K9me3 and H3K27me3. This H3K9me2-enriched state is often collapsed with the H1-rich chromatin of the A compartment as ‘naïve chromatin’, yet both show functional differences with regard to their composition and regulation, notably the requirement on cell division for opening of lamin-associated loci.

Strikingly, PAX7 action in the A and B compartments is associated with functionally distinct biological outputs. PAX7-regulated genes in the A compartment are broadly expressed and involved in general cellular functions; PAX7-regulated genes in the B compartment encode cell-type-specific factors central to melanotrope identity, and these are activated de novo from previously silent states. Cross-tissue analysis of public H3K4me1 and H3K27ac data shows that B-compartment PAX7-pioneered enhancers are far more cell-type-restricted than their A-compartment counterparts. This functional partition is consistent with the broader role of B compartments and lamina-associated domains in sequestering lineage-specific genes for repression in inappropriate cell types (*37, 38*). Cell-fate specification by PAX7 therefore involves not just the activation of new genes but the spatial reorganization of those genes from a sequestered, lamina-associated state into the active nuclear interior, a feature that may distinguish bona fide developmental specification from transcriptional fine-tuning of an already-established programme.

In summary, PAX7 acts through two compartment-defined pioneer modes that differ in mechanism, kinetics, scale, and biological function. By coupling local enhancer activation to domain-scale compartment switching, the B-compartment mode provides the spatial reorganization required for de novo activation of cell-type-specific genes and thereby implements the developmental component of cell-fate specification. The A-compartment mode modulates the broader transcriptional context in which that programme is expressed; it is noteworthy that genes and enhancers that acquire transcriptional memory are primarily within this group (Harris et al. In submission). Together, the two modes deliver the full transcriptional and epigenetic programme that defines a new cell identity.

## Materials and Methods

### ChIP-seq and ATAC-seq processing

Paired-end reads were trimmed for adapter content and aligned to the mouse mm10 reference genome using Bowtie (*39*) 2.5 (--no-unal --no-mixed --no-discordant). Mapping results were processed using Samtools (*40*) 1.22 to fix mate-pair information (fixmate), coordinate-sort (sort) and remove duplicates (markdup -r). Aligned reads were piled up, genomic coverage normalized to counts per millions at a resolution of 10 bp, and bigWig files were created using wigToBigWig (*41*) from UCSC Genome Browser Utilities v486. Peaks were called using callpeak (-f BAMPE -p 1e-3/1e-5 -g mm --min-length 100 --max-gap 50) from MACS (*42*) 2.2.9.1.

### ChIP-seq and ATAC-seq analyses

Local signal was quantified by the mean over a 500 bp window (1,000 bp for H3K4me1, H3K4me3 and H3K27ac; 4,000 bp for repressive histone marks, histone H1/H3 and lamin A/C) or a specified window. ChIP-seq and ATAC-seq signals were normalized relative to input signal, and relative to the average signals at random loci and constitutively active enhancers. PAX7-bound loci with low/high input signal (outside the first/third quartile minus/plus half the interquartile range of the signal for all bound loci), localized at promoters (within a 1,000 bp window of a gene’s TSS (GENCODE M25) or exhibiting a H3K4me3 signal before *Pax7* expression greater than the third quartile plus 1.5 times the interquartile range of the signal for all bound loci) or localized at insulator (exhibiting a CTCF signal before *Pax7* expression greater than the third quartile plus 1.5 times the interquartile range of the signal for all bound loci) were excluded (table S1). PAX7-bound loci were categorized considering the H3K4me1 deposition and DNA accessibility (ATAC) before and after stable *Pax7* expression, as shown in fig. S1B.

### Micro-C processing

Paired-end reads were aligned to the mouse mm10 reference genome using BWA-MEM2 (*43*) 2.2.1 (-5 -S -P -T 0). Mapping results were parsed, sorted, and deduplicated using Pairtools (*44*) 1.1.3, filtering for high-quality alignments (--min-mapq 40) and specific ligation events (--walks-policy 5unique --max-inter-align-gap 30). Pairs were indexed using Pairix (*45*) 0.3.7, and corresponding bam files were coordinate-sorted and indexed using Samtools (*40*) 1.22. Contact matrices were generated in hic format using pre from Juicer (*46*) 1.8.0, including raw, Knight-Ruiz (KR) normalized and observed-over-expected signal. Topologically associating domains (TAD) were called using OnTAD (*47*) 1.4 (-minsz 3 -maxsz 200 -lsize 5 -ldiff 1.96 -penalty 0.1) at a resolution of 5,000 bp (table S3).

### Micro-C analyses

TAD-level analyses were performed at TADs between 100 kb and 1 Mb harbouring at least one Pioneer enhancer or at least one Primed enhancer and no Pioneer enhancer. If applicable and in the case of nested TADs, one TAD was assigned per enhancer, selecting the TAD with the largest difference in compartment-score variation relative to its 200 kb flanking regions. Aggregate analyses were performed on KR-normalized contact matrices at 5,000 bp resolution, spanning the TAD extended by flanking regions equal to its length on either side. Matrices were linearly interpolated onto a common 200*200 grid and averaged across TADs before and after stable *Pax7* expression. Insulation scores were computed on the size-normalized maps as the log2 mean contact frequency within a sliding off-diagonal (diamond) window of 18 rescaled bins, relative to the map-wide average, and boundary insulation was taken at the rescaled 5′ and 3′ TAD boundaries.

### Genomic compartments analyses

Compartment scores were computed from intra-chromosome KR-normalized observed-over-expected contact matrices at 5,000, 10,000 and 100,000 bp resolutions. For each chromosome, values were clipped to the third quartile plus three times the interquartile range and eigendecomposition was performed to extract the three leading eigenvectors. Eigenvectors whose positive and negative entries were strongly imbalanced (difference exceeding 80% of the vector length) were excluded, and the remaining eigenvectors were selected and oriented by their correlation with lamin A/C ChIP-seq signal binned at the corresponding resolution. Compartment scores were computed as the product of the selected eigenvector and the square root of its eigenvalue divided by the vector length. For cross-sample comparisons, quantile normalization was applied to compartment scores from individual samples using compartment scores computed from merged contact matrices as reference.

### Sub-compartments analyses

Principal component analysis was performed on a KR-normalized matrix composed of trans contacts scores between even and odd chromosomes at a resolution of 100 kb, thereby isolating the compartment signal from cis-contact bias. K-means clustering of the average H3K4me1, H3K4me3, H1, H3K9me2, lamin A/C, H3K9me3 and H3K27me3 signal over the 100 kb bins was performed to associate each bin to a sub-compartment (table S2). The cluster count was chosen empirically as the maximum number of clusters not resulting in duplicated clusters (i.e., with similar chromatin landscape).

### RNA-seq processing

Paired-end reads were trimmed for adapter content and aligned to the mouse mm10 reference genome using STAR (*48*) 2.7.11, generating both coordinate-sorted and transcriptome-projected BAM files (--quantMode TranscriptomeSAM) based on GENCODE (*49*) M25. Gene expression was quantified using RSEM (*50*) 1.3.3 (--paired-end --alignments) from the transcriptome-aligned reads.

### RNA-seq and gene ontology analyses

Gene expression values (transcripts per million, TPM) were normalized across samples using the median-of-ratios method (*51*): for each gene with non-zero expression in all samples, a pseudo-reference was defined as the geometric mean of its TPM values across samples, and per-sample normalization factors were obtained as the median ratio of the pseudo-reference to the TPM values across all genes. Genes regulated by PAX7 are selected as shown in Fig. 5A, and their A/B compartment association is determined by the compartment score at their TSS with a resolution of 10,000 bp. Gene ontology was performed on genes associated with the A and B compartment separately, and either regulated, downregulated or upregulated by PAX7, using PANTHER (*52*) 19.0 overrepresentation test with the GO biological process annotation set (table S4).

### ChIP-Atlas analyses

ChIP-seq peak-calling data from ChIP-Atlas (*28*), encompassing 2,008 H3K4me1 and 4,536 H3K27ac samples across 237 and 368 cell lines or tissues, respectively, were downloaded (table S4). Each sample was overlapped to PAX7-bound loci in our system and grouped by cell line or tissue of origin (as catalogued). A PAX7-bound locus was considered positive in another cell line if it colocalized with at least one peak from a sample in that cell line.

### Statistics and reproducibility

Statistical tests are described in the figure legends. Genome-wide sequencing samples, replicates and parameters are listed in table S6. Micro-c experiments are duplicates before *Pax7* expression and singlets after stable *Pax7* expression. They were validated against multiple ChIP-seq chromatin markers, as exemplified in Fig. S2C, Fig. 2 and Fig. 4A-E. RNA-seq are duplicates. ChIP-seq and ATAC-seq are either duplicates or singlets with at least two equivalent and concordant ChIP-qPCR or ATAC-qPCR biological replicates. Antibodies are listed in table S6.

## Acknowledgments

We are grateful to Sarah Boissel for next-generation sequencing and to Valerie Magoon for expert secretarial assistance.

## Funding

Canadian Institutes of Health Research grant FDN-154297 (JD)

Digital Research Alliance of Canada computation allocation zmv-553 (JD)

## Author contributions

Conceptualization: AG, JD

Investigation: AG, JH, VLR

Data Curation: AG, AM, AB

Formal Analysis: AG

Supervision: JD

Writing – original draft: AG, JD

## Competing interests

Authors declare that they have no competing interests.

## Data, code, and materials availability

Sequencing data are available on GEO under GSE87185, GSE125671, GSE173743, GSE225231, GSE240090 and GSE335231. Per-sample accession IDs are provided in table S6.

## Supplementary Materials

Figs. S1 to S5

Tables S1 to S6

## Notes

### Competing Interest Statement

The authors have declared no competing interest.

