## Supplementary Figures for "Chromatin compartments define distinct modes of action and biological roles of the pioneer Pax7"

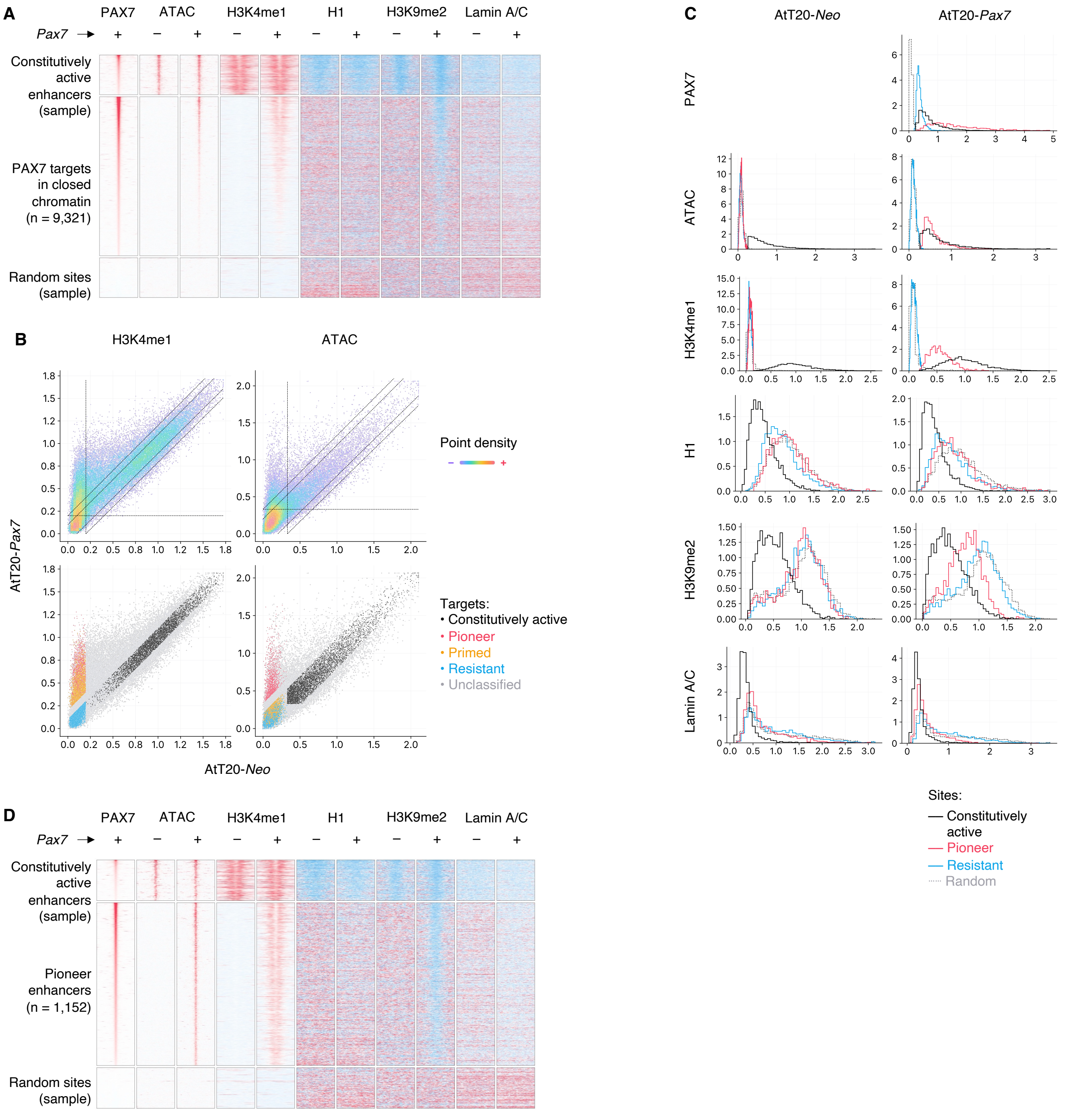


**Fig. S1. Genome-wide binding of PAX7.** (**A**) Heatmaps of the indicated markers at multiple subsets of PAX7-targeted sites with or without stable *Pax7* expression. Constitutive and random sites subsampled. Sites ordered by PAX7 recruitment. 4 kb window centered on PAX7 recruitment. (**B**) H3K4me1 deposition (ChIP-seq) and DNA accessibility (ATAC-seq) at PAX7 binding sites with or without *Pax7* expression. Bottom plots show selection threshold for constitutively active (n = 3,984), pioneered (n = 1,152), primed (n = 2,523) and resistant (n = 2,761) sites bound by PAX7 (total n = 28,504). (**C**) Distribution of the indicated markers signal at multiple subsets of PAX7-targeted with or without stable *Pax7* expression. (**D**) Heatmaps of the indicated markers at multiple subsets of PAX7-targeted sites with or without stable *Pax7* expression. Constitutive and random sites subsampled. Sites ordered by PAX7 recruitment. 4 kb window centered on PAX7 recruitment.

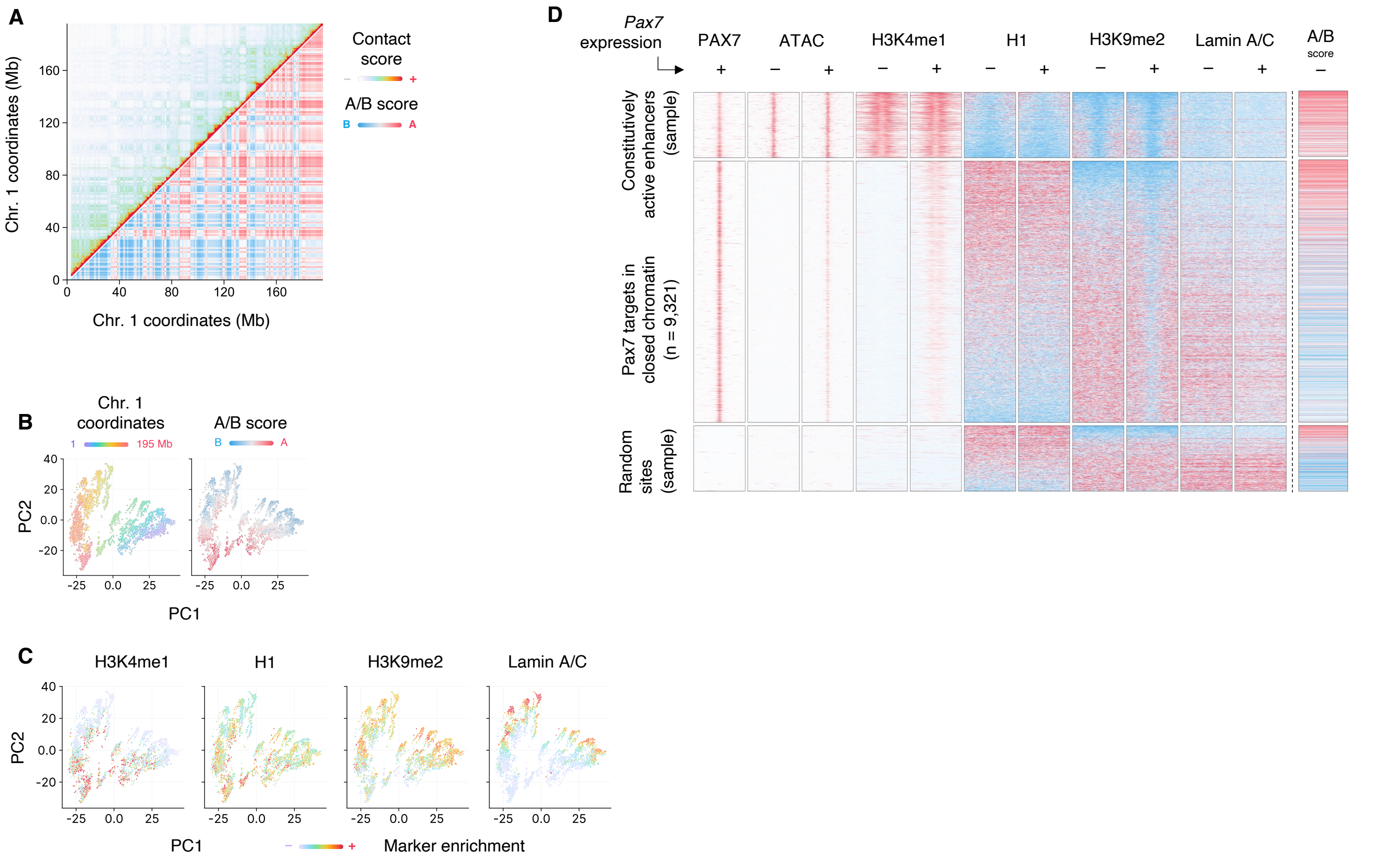


**Fig. S2. Genome compartments and chromatin states.** (**A**) Micro-C contact map at resolution of 100 kb of AtT20 chromosome 1 after *Pax7* expression (top left). Corresponding A/B compartment identity (bottom right). (**B**) PCA of Micro-C contacts without *Pax7* expression of AtT20 chromosome 1 at a resolution of 10 kb, with bin coordinate or A/B score color-coded. (**C**) Average value of the indicated marker per bin from (B). (**D**) Heatmaps of the indicated markers at multiple subsets of PAX7-targeted sites with or without stable *Pax7* expression. Constitutive and random sites subsampled. Sites ordered by the ratio of initial H1 over H3K9me2 signal. 4 kb window centered on PAX7 recruitment.

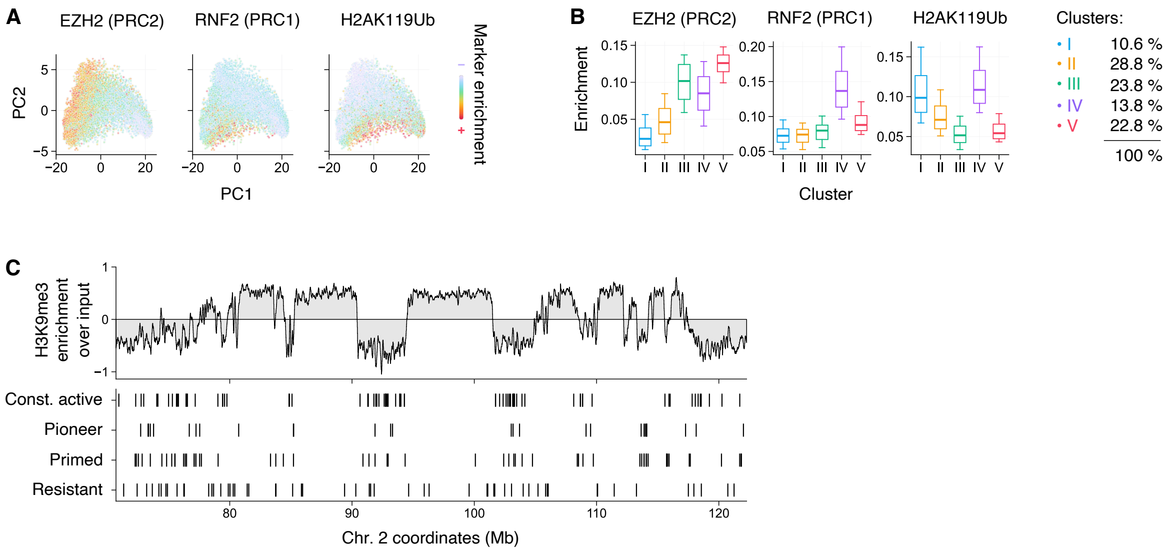


**Fig. S3. Chromatin state at sub-compartments.** (**A**) Average value of the indicated marker per bin from the PCA from Fig. 2B. (**B**) Quantification of the specified markers without *Pax7* expression at the clusters from Fig. 2D. All pairwise comparisons are statistically significant. (**C**) H3K9me3 ChIP-seq and PAX7 targets sites by category at a representative genomic region. P-values computed from two-sided Mann-Whitney U tests; boxplots represent the 10-25-50-75-90$^{th}$ percentiles.

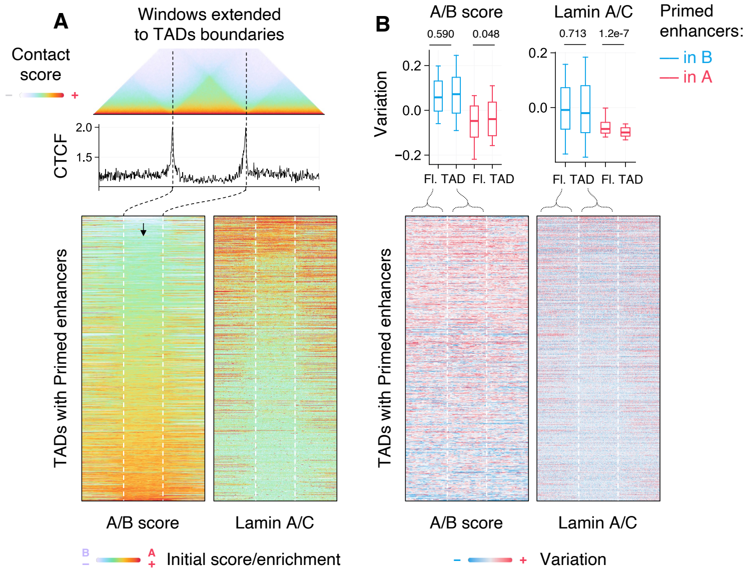


**Fig. S4. Domain-wide effects of PAX7.** (**A**) Average Micro-C contact map and average CTCF ChIP-seq profile before *Pax7* expression at unique and unnested TADs harboring Primed enhancers and no Pioneer enhancers (top). Heatmaps of initial A/B score and lamin A/C association at these TADs, ordered by average A/B score (bottom). (**B**) Heatmaps of A/B score and lamin A/C association variation before and after *Pax7* stable expression at TADs from (A). Distribution of the average variation within the TADs or the 200 kb flanking regions (Fl.) at the ¼ Primed enhancers with strongest B or A association is indicated on top. P-values computed from two-sided Mann-Whitney U tests; boxplots represent the 10-25-50-75-90th percentiles.

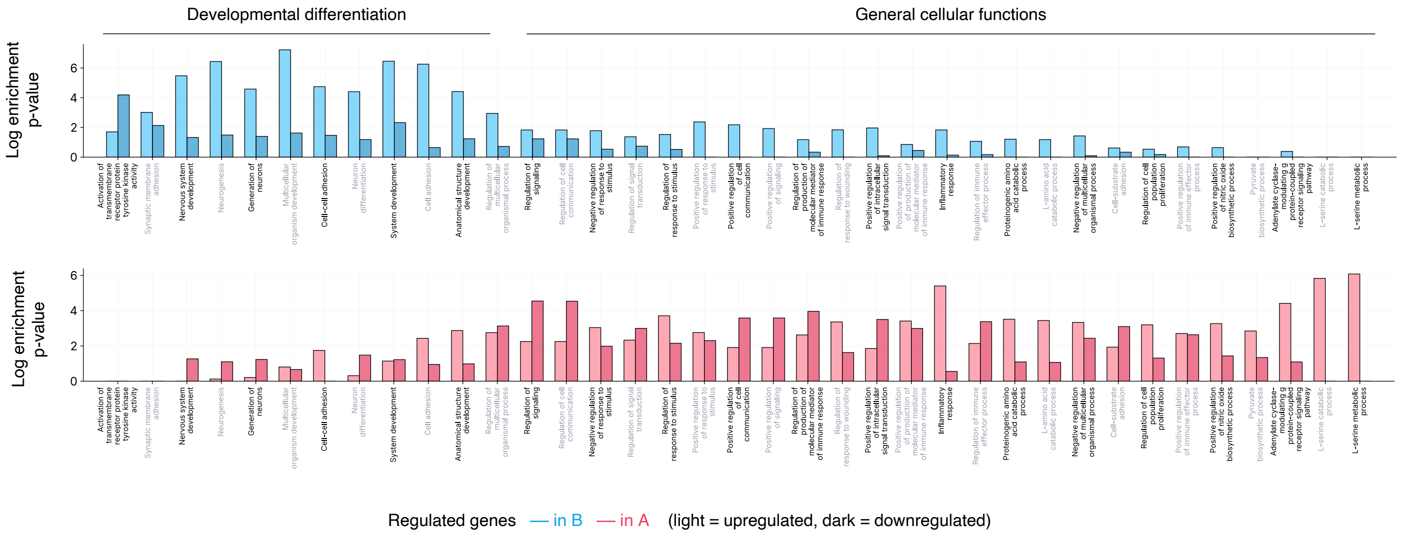


**Fig. S5. Transcriptional action of PAX7.** Gene ontology (GO, biological process) analysis of PAX7-regulated genes in A vs. B compartment. Terms ordered by the difference in A vs. B enrichment. Grayed terms for visualization purpose only.
